# Unexpected activity of oral fosfomycin against resistant strains of *Escherichia coli* in murine pyelonephritis

**DOI:** 10.1101/627984

**Authors:** Annabelle Pourbaix, François Guérin, Charles Burdet, Laurent Massias, Françoise Chau, Vincent Cattoir, Bruno Fantin

## Abstract

Fosfomycin-tromethamine activity is well established for oral treatment of uncomplicated lower urinary tract infections but little is known about its potential efficacy in pyelonephritis. Ascending pyelonephritis was induced in mice infected with 6 strains of *Escherichia coli* (fosfomycin MICs: 1 μg/ml to 256 μg/ml). Urine pH was 4.5 before infection and 5.5-6.0 during infection. Animals were treated for 24h with fosfomycin (100 mg/kg subcutaneously every 4 hours) and CFU were enumerated in kidneys 24h after the last fosfomycin injection. Peak (20.5 μg/ml at 1h) and trough (3.5 μg/ml at 4h) levels in plasma were comparable to those obtained in human after an oral dose of 3 grams. Fosfomycin treatment significantly reduced bacterial loads in kidneys (3.65 log_10_CFU/g [min-max=1.83-7.03] and 1.88 log_10_CFU/g [1.78-5.74] in start-of-treatment control mice and treated mice, respectively, *P* < 10^-6^). However, this effect was not found to differ across the 6 study strains (P = 0.71) and between the 3 susceptible and the 3 resistant strains (P=0.09). Three phenomena may contribute to explain this unexpected *in vivo* activity: i) in mice, fosfomycin kidney/plasma concentrations ratio increased from 1 to 7.8 (95% CI, 5.2; 10.4) within 24 hours; *in vitro*, when pH decreased to 5: (ii) fosfomycin MICs for the 3 resistant strains (64-256 μg/ml) decreased into the susceptible range (16-32 μg/ml) and: iii) maximal growth rates significantly decreased for all strains and were the lowest in urine. These results suggest that local fosfomycin concentrations and physiological conditions may favour fosfomycin activity in pyelonephritis, even against resistant strains.

## INTRODUCTION

Over the last two decades, resistance to β-lactams among *Enterobacteriaceae* has emerged as a major public-health threat. Isolates of *Escherichia coli* producing extended-spectrum β-lactamases are currently responsible for a large proportion of urinary tract infections (UTIs), in the community as well as in the healthcare setting (1–3). In the current era of increasing prevalence of antibiotic resistance, fosfomycin has attracted renewed interest for the treatment of infections caused by multidrug-resistant pathogens, and especially UTIs. Indeed, it has a broad-spectrum antimicrobial activity and a favorable safety profile (4, 5). Fosfomycin-tromethamine, a soluble salt with improved bioavailability over fosfomycin, is currently recommended in single-dose as the first-line drug for the treatment of uncomplicated lower UTIs in Europe (6) and in the United States (7). However, it is unknown whether fosfomycin-tromethamine would be useful for the treatment of pyelonephritis in human, raising the questions of kidney diffusion and breakpoints to be used in pyelonephritis as compared with cystitis. Indeed, the current susceptibility breakpoint of fosfomycin for *Enterobacteriaceae* is a minimum inhibitory concentration (MIC) of 32 μg/ml according to the European Committee on Antimicrobial Susceptibility Testing (EUCAST) (8). We previously demonstrated that fosfomycin resistance in *E. coli* strains from various genetic backgrounds was associated with a decrease of *in vitro* fitness and *in vivo* virulence in a murine model of pyelonephritis (9). In addition, some mutations conferring fosfomycin resistance have been shown to decrease pilus biosynthesis and bacterial adhesion to epithelium cells (10, 11). Furthermore, Martin-Gutierrez et al. recently demonstrated that anaerobiosis and acidic pH values in urine decreased fosfomycin MICs in strains harboring chromosomal resistance mutations (12). Finally, it has been previously shown that high concentrations (1000-4000 μg/ml) were achieved in urine after oral administration of 3 g of fosfomycin-tromethamine in humans and remained above 100 μg/ml for at least 30-40h (13), but data on concentrations in kidneys are scarce.

However, possible limitations for the use of the oral fosfomycin-tromethamine in pyelonephritis may be anticipated, related to potentially low concentrations in kidneys as compared to urine, which in turn might be associated with limited bactericidal activity and risk of selection of resistant mutants. Indeed, the selection of spontaneous fosfomycin-resistant mutants occurs at a very high rate *in vitro* (between 10^-7^ to 10^-6^ cells among Gram-negatives) (14).

Therefore, the aim of the present study was to investigate the activity of fosfomycin in a murine model of pyelonephritis due to *E. coli,* using a dosing regimen that reproduced plasma concentrations comparable to those obtained in human with the oral formulation of fosfomycin-tromethamine. Different strains with increasing levels of MICs were used in order to define the *in vivo* activity of fosfomycin-tromethamine according to strain susceptibility in this specific infection.

## RESULTS

### Bacterial study strains

Characteristics of bacterial strains used in the study are shown in Table 1. Three strains were susceptible to fosfomycin and three strains were resistant, according to EUCAST breakpoints (32 μg/ml). Mutations in genes known to be involved in fosfomycin resistance (*uhpB, uhpC, glpT* and *cyaA*) were detected in all strains, including those categorized as susceptible (B175 and B56) according to the current EUCAST breakpoint (Table 1). All but one strain (*E. coli* B56, producing an extended spectrum β-lactamase) were susceptible to other antibiotic families.

**Table 1.** Susceptibility to fosfomycin of *E. coli* strains used in this study, according to pH and media.

| Strains | CFT073 | B56 | B175 | MUT2 | C05 | C114 |
| --- | --- | --- | --- | --- | --- | --- |
| Origin | Urine clinical isolate | Urine clinical isolate | Urine clinical isolate | Mutant from CFT073 | Urine clinical isolate | Urine clinical isolate |
| Phylogenetic group | B2 | B2 | B2 | B2 | B2 | B2 |
| Fosfomycin MIC* according to pH: |  |  |  |  |  |  |
| pH 7.2 | 1 | 8 | 32 | 64 | 128 | 256 |
| pH 6 | 1 | 8 | 32 | 16 | 64 | 32 |
| pH 5 | 1 | 16 | 32 | 16 | 32 | 32 |
| Susceptibility categorization according to pH 7.2/pH 5** | S / S | S / S | S / S | R / S | R / S | R / S |
| Detected mutation(s) of fosfomycin resistance | None | UhpB (P169S) | CyaA (L125F),<br>UhpC (T72I) | UhpB (G469R) | UhpC (Q210X) | UhpC (Q132X),<br>GlpT (G144D) |

\*: MICs are in µg/ml
\*\*: S denotes susceptible and R resistant, according to EUCAST breakpoints <sup>7</sup>

### Fosfomycin concentrations in plasma and kidneys

Fosfomycin concentrations in plasma and kidneys after a single injection of 100 mg/kg sc are shown in Table 2. While fosfomycin concentrations were comparable in plasma and kidneys 1h following the injection, fosfomycin concentrations were higher in kidneys than in plasma over time, with mean concentrations at 24h < 1 mg/L in plasma and 6.8 mg/L in kidneys, and a kidney/plasma ratio of 7.8 (95% CI, 5.2; 10.4) (Figure 1). After a single injection, area under the concentrationtime curve in kidney (AUC_0-24h_) was estimated to 166.6 mg.h/L, being two-fold higher than in plasma where it reached 85.0 mg.h/L. Fosfomycin AUC_0-24h_ in plasma and kidneys after a 100 mg/kg injection every 4h for 24h were 357.1 and 614.5 mg.h/L, respectively and AUC in plasma to the time a sacrifice (AUC_0-44h_) was 470.3 mg.h/L.

**Figure 1.**
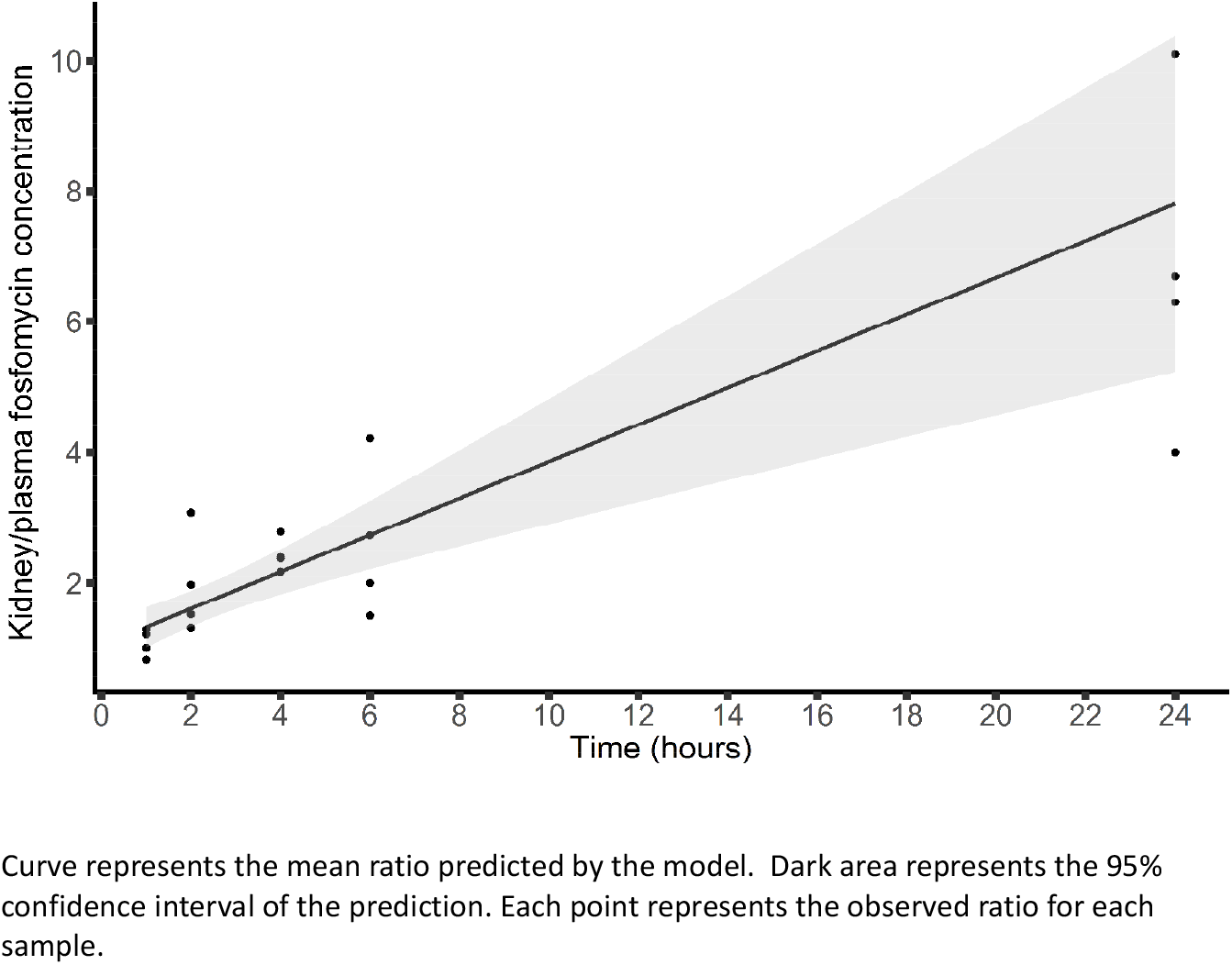
Ratio of the kidney/plasma fosfomycin concentrations in CBA mice after a single injection of 100 mg/kg sc.

**Table 2.** Fosfomycin concentrations in plasma and kidneys from CBA female mice after a single injection of 100 mg/kg given subcutaneously. Each set of values for a given sample time corresponds to the mean of 4 mice (min-max).

| Time | Fosfomycin concentrations (µg/ml) in: |  |
| --- | --- | --- |
|  | Plasma | Kidneys |
| 1h | 20.5 (9.0 – 33.5) | 20.8 (11.4 – 27. 5) |
| 4h | 3.5 (2.8 – 4.8) | 8.4 (6.7 – 11.4) |
| 24h | < 1 (<1 - < 1) | 6.8 (4.0 – 10.1) |

### Fosfomycin antimicrobial effect in murine pyelonephritis

The bacterial loads in kidneys according to fosfomycin treatment and the infective strain are presented in Figure 2 and Table 3. Fosfomycin treatment significantly reduced the bacterial loads in kidney (*P* < 10^-6^). Median (min-max) bacterial loads in kidney were 3.7 log10 CFU/g (1.8-7.0) in start-of-treatment control mice, and 1.9 log10 CFU/g (1.8-5.7) in fosfomycin-treated mice. However, there was no significant difference in kidney bacterial loads according to the infective strain (*P* = 0.71) and between the 3 susceptible and the 3 resistant strains (*P* = 0.09). The interaction between the infective strain and the fosfomycin effect was not significant (*P* = 0. 53). Similarly, the proportion of sterile kidneys in treated mice did not differ between the 6 strains (P > 0.8) (Table 3). No fosfomycin-resistant mutant was detected in kidneys at the time of sacrifice for any of the 6 strains.

**Figure 2.**
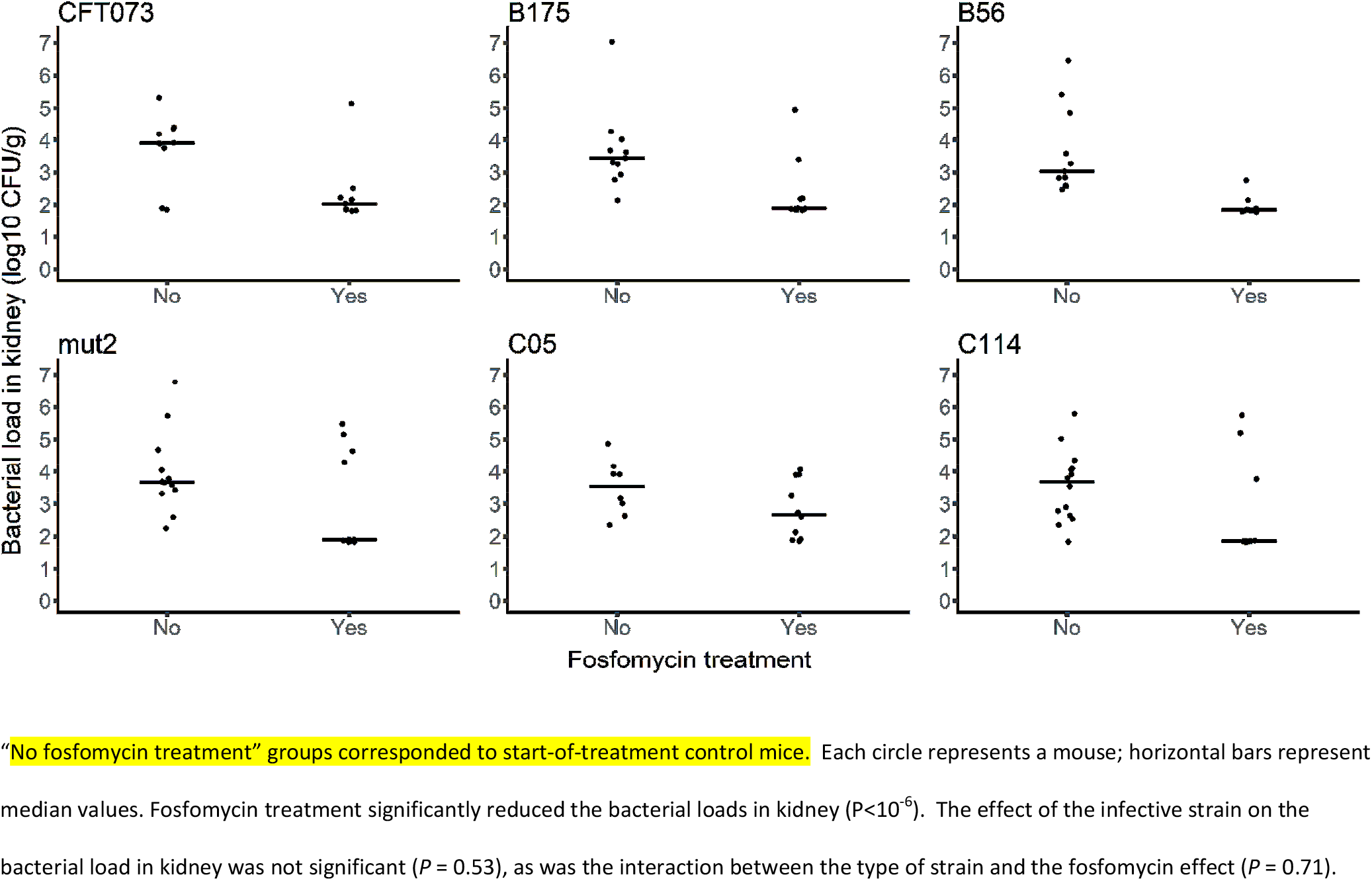
Bacterial counts (log_10_ CFU per gram of kidney) of *E. coli* in mice with pyelonephritis treated or not for 24h with fosfomycin 100 mg/kg sc every 4 hours, according to study strains.

**Table 3.** Pharmacokinetics/pharmacodynamics of fosfomycin (FOS) in murine pyelonephritis due to *E. coli* after a 24h treatment.

| Strains | FOS MIC<br>(µg/ml) | FOS AUC<sub>0-24</sub><br>plasma/MIC | FOS AUC<sub>0-24</sub><br>kidney/MIC | Median log<sub>10</sub> CFU counts/g of kidneys<br>[range min-max] |  |
| --- | --- | --- | --- | --- | --- |
|  |  |  |  | Start-of-treatment control mice | FOS treated mice |
|  |  |  |  | (sterile/ total) | (sterile/ total) |
| CFT073 | 1 | 357.1 | 614.5 | 3.9 [1.9-5.3] (0/11) | 2.0 [1.8-5.1] (4/9) |
| B56 | 8 | 44.6 | 76.8 | 3.0 [2.5-6.5] (0/11) | 1.8 [1.8-2.8] (4/10) |
| B175 | 32 | 11.2 | 19.2 | 3.4 [2.1-7.0] (0/11) | 1.9 [1.8-4.9] (4/10) |
| MUT2 | 64 | 5.6 | 9.6 | 3.7 [2.2-6.8] (0/12) | 1.9 [1.8-5.5] (5/10) |
| C05 | 128 | 2.8 | 4.8 | 3.7 [2.3-4.9] (0/8) | 2.7 [1.9-4.1] (3/10) |
| C114 | 256 | 1.4 | 2.4 | 3.7 [1.8-5.8] (0/14) | 1.9 [1.8-5.7] (6/10) |

CFU: colony-forming units, FOS: fosfomycin, MIC: minimum inhibitory concentration. Median (min-max) bacterial loads in kidney were 3.7 log<sub>10</sub> CFU/g (1.8; 7.0) in start-of-treatment control mice, and 1.9 log<sub>10</sub> CFU/g (1.8; 5.7) in fosfomycin-treated mice ( $P < 10^{-6}$ ). Lower limit of detection was 1.8 log<sub>10</sub> CFU/g of kidney.

### Pharmacokinetic/pharmacodynamic (PK/PD) analysis

Fosfomycin AUC_0-24_/MIC ratios in plasma and kidneys according to strains are shown in Table 3. The relationship between the bacterial loads in kidneys and the AUC_0-24_/MIC ratios in plasma or kidney and the AUC_0-44_/MIC ratios in plasma were not significant (P=0.45 for both plasma and kidney).

### Effect of pH on fosfomycin activity and bacterial growth rate

In order to explain the unexpected activity of fosfomycin against both susceptible and resistant strains, investigations were performed to test the influence of acidic pH, as 85% of patients with UTI due to *E. coli* had a urine pH of < 6.5) (12). Indeed, pH in uninfected urine from 5 CBA mice before experimental pyelonephritis was 4.5 and ranged from 5.5 to 6.0 in the same mice after 48h of experimental pyelonephritis.

### Effect of pH on in vitro fosfomycin activity

Low pH values increased fosfomycin activity among fosfomycin-resistant strains (Table 1). Indeed, when pH decreased from 7.2 to 5, fosfomycin MICs against the 3 resistant strains decreased from 64-256 μg/ml to 16-32 μg/ml, which corresponded to susceptibility according to EUCAST (8).

### *Effect of pH and urine on* in vitro *bacterial growth rate*

At pH 7, maximal growth rates (MGRs) differed among the 6 studied strains, with median values ranging from 3.29 to 4.20 h^-1^ (*P* < 0.01). For each of the 6 strains, MGR was reduced as the pH was lower (Figure 3). This reduction was statistically significant (*P* < 0.05) between pH 7 and pH 5 for all strains except C05. The lowest MGR values were observed in urine with a significant reduction as compared with MGR value determined at pH 7 for all strains (*P* < 0.05). For each of the 6 strains, time to achieve MGR was prolonged at pH 5 as compared with pH 7 and this difference was statistically significant (*P* < 0.05) for all strains except C05 (Figure 3).

**Figure 3.**
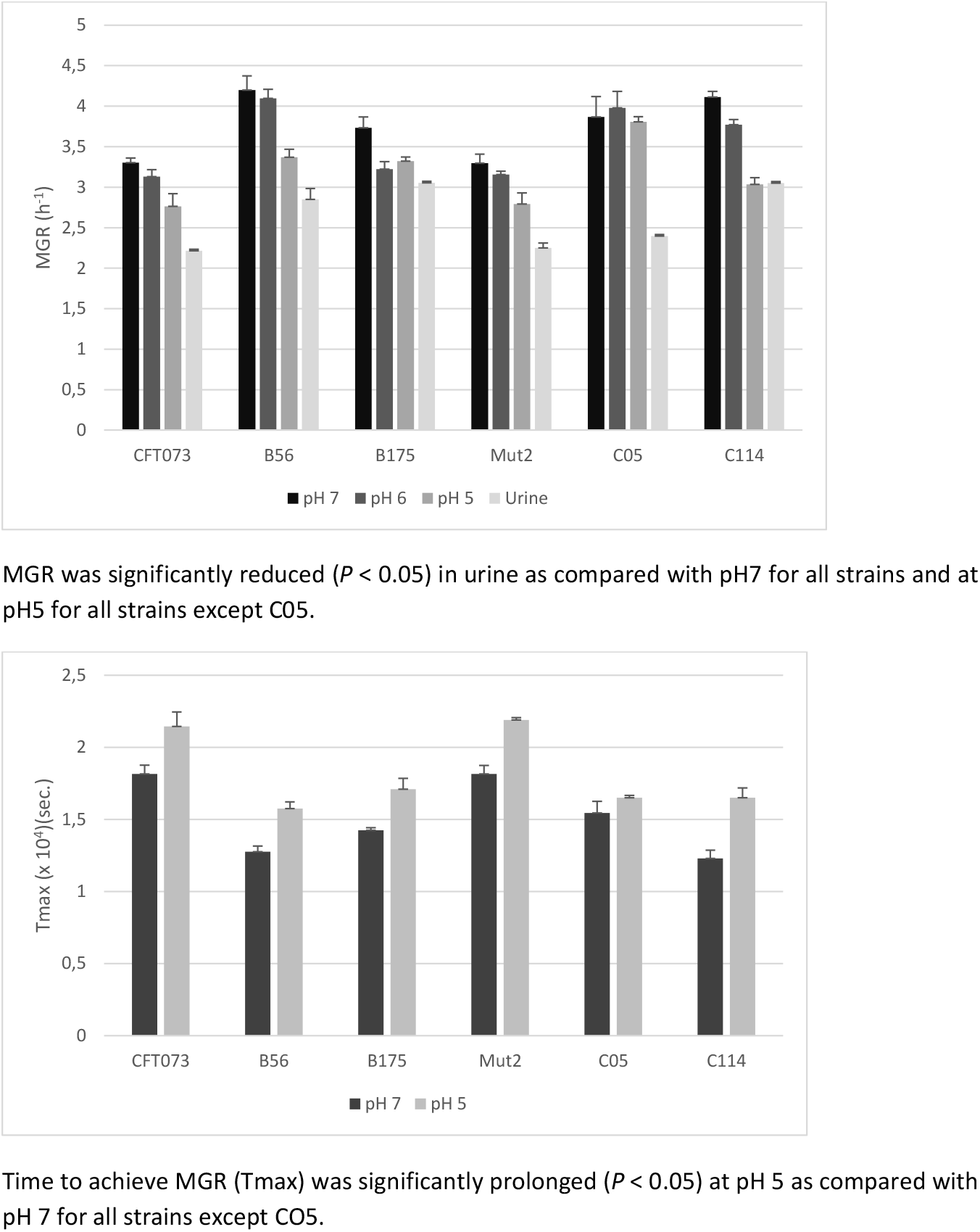
In vitro maximal growth rates (upper panel) and time to achieve maximal growth rate (lower panel) for each study strain according to pH in Luciani-Bertani or in urine. Each value is the median of three independent experiments. Brackets represent standard deviations.

## DISCUSSION

Fosfomycin has been approved for decades as an injectable antibiotic for the treatment of systemic infections due to both Gram-positive cocci and Gram-negative bacilli, including severe infections (4, 15, 16). More recently, an oral formulation, fosfomycin-tromethamine, has rapidly become the first-line empirical treatment recommended worldwide as a single dose for acute uncomplicated lower UTIs (6, 7).

Here we show that, when given a regimen generating peak and trough levels in plasma in the range of those obtained in human after a single dose of the oral formulation of fosfomycin-tromethamine (approximatively 20 μg/ml and 5 μg/ml, respectively) (17, 18), fosfomycin produced a significant reduction of bacterial load after 24h of treatment in kidneys from mice with pyelonephritis. However, the unexpected result was that fosfomycin efficacy was similar whatever the study strains, susceptible or resistant to fosfomycin, according to EUCAST (8).

Fosfomycin activity observed in infected kidneys may result from the combination of local specific physiological conditions in urine and of fosfomycin concentrations in the kidneys. Indeed, several authors have already shown that low pH values increased fosfomycin *in vitro* activity (12, 19, 20). In an acidic environment such as urine, fosfomycin is partially protonated, being in a more lipophilic state, allowing fosfomycin entry into bacteria, and resulting in higher antimicrobial activity (20). Our results support these findings, as low pH values were associated with a significant decrease in fosfomycin MIC values for resistant strains into the susceptible range (Table 1). Other factors than acidic pH may alter the growth of *E. coli* in urine, such as the lack of essential sources of metabolic elements like urea, creatinine, iron, citric acid, D-serine, or ammonia (21).

In the present study, we also confirmed *in vitro* that low pH values were associated with a decreased *in vitro* fitness for fosfomycin-susceptible or resistant strains, as previously reported (12) and showed that this decrease was maximal in urine for all the tested strains (Figure 2).

Chromosomal mutations conferring fosfomycin resistance have been associated with a high biological cost, entailing a reduced fitness (11). This fitness cost is of particular interest in uropathogenic *E. coli* strains because if the biological cost is high, the resistant bacteria will not grow at the minimal rate needed to establish infection or to invade the kidney (22, 23). We have recently shown in the same model of ascending murine pyelonephritis due to uropathogenic *E. coli* strains belonging to the B2 phylogenetic group that there was a significant reduction in kidney infection rates with fosfomycin-resistant isolates as compared with susceptible ones (9). This phenomenon may of course have favoured fosfomycin antimicrobial activity in mouse pyelonephritis and have contributed to limit the selection of fosfomycin-resistant mutants, as observed in the present study in mice with no detection of fosfomycin resistance for any strain tested.

On a pharmacokinetic point of view, fosfomycin concentrations over time were higher in kidneys than in plasma. This increase in drug exposure obviously favored fosfomycin activity in pyelonephritis. On a PK/PD point of view, recent studies done in mice demonstrated that the PK/PD index that best predicted fosfomycin efficacy in a thigh and in UTI model was AUC/MIC ratio (24, 25). More specifically, in the thigh infection model, for *E. coli,* net stasis was observed at a median AUC/MIC ratio value of 19.3 and one-log kill was observed at a median AUC/MIC ratio value of 97.5 (24). According to these data, since a similar dosing regimen was used in the present study in all mice and generated in plasma the same AUC_0-44h_ of 470.3 mg.h/L at the time of sacrifice, the target value for net stasis would be achieved only for strains with an MIC of 24 μg/ml or lower, and the one-log kill target would be achieved only for strains with an MIC of 4 μg/ml or lower. The “higher than expected” fosfomycin activity observed in our model (Table 3) is probably the consequence of the overall favorable local physiological conditions discussed above, reducing the MIC, fitness and virulence of resistant strains and associated with kidney concentrations favouring fosfomycin activity in pyelonephritis.

The lack of correlation we observed between AUC/MIC ratio in plasma and CFUs in kidneys was related to the fact that, on one hand, all the mice were treated with the same dosing regimen and therefore exposed to the same AUC, and on the other hand, local MICs of resistant strains in urine became very similar to those from susceptible strains. Thus, as AUC was similar for all strains and MICs in local conditions not as different as in standard conditions (Table 1), the AUC/MIC ratio reached a “plateau” that precluded to investigate the PKPD relationship and did not allow us to determine an MIC breakpoint.

Several limitations of our study, due to experimental conditions, should be taken into account when extrapolating the data as they could limit the probability for selection of fosfomycin-resistant mutants *in vivo:* i) bacterial loads in kidneys before fosfomycin treatment were moderate, and no spontaneous mortality is observed in this model; ii) duration of fosfomycin treatment was limited to 24h, a short duration of time to select for resistant mutants; however, it must be acknowledge that this is very similar to the situation in human after an oral single dose of fosfomycin; iii) the fosfomycin dosing regimen used in the present study generated an AUC_0-24h_ that corresponded approximately to what would be obtained in human after two doses of oral fosfomycin-tromethamine (15, 16, 18), as repeated oral doses is a dosing regimen that is under investigation for other situations than cystitis in women (19); iv) finally, we did not perform a group of end-of-treatment control mice which would have help to analyze the relative part of spontaneous bacterial clearance from the kidneys and the antibacterial effect of fosfomycin.

In conclusion, our results suggest that fosfomycin-tromethamine oral formulation might be of interest in human for the treatment of UTI with parenchymal infections, such as pyelonephritis, due to favorable local physiological conditions and kidney concentrations. Our results suggest that repeated dosing of fosfomycin-tromethamine should be investigated for the treatment of uncomplicated pyelonephritis in women.

## MATERIALS AND METHODS

### Bacterial strains

Six bacterial strains of *E. coli* were used (Table 1), with MICs of fosfomycin ranging from the susceptible to resistant range (1 to 256 μg/ml). The reference wild-type *E. coli* CFT073 (O6:K2:H1) strain (25) was previously used to set up a murine model of pyelonephritis by our group (26, 27). Other strains were clinical isolates from urinary tract infections (UTIs), except *E. coli* MUT2 that was a fosfomycin-resistant mutant selected *in vitro* from *E. coli* CFT073. We selected strains belonging to phylogenetic group B2 since such strains are most frequently responsible for UTIs in humans and carry the most important virulence factors (28, 29).

### *In vitro* fosfomycin activity

MICs of fosfomycin (Sanofi-Aventis, Paris, France) were determined by the dilution method in Mueller-Hinton agar (pH 7.2) in accordance with EUCAST guidelines (9), with 25 μg/ml glucose-6-phosphate (G6P) (Sigma–Aldrich, Saint-Quentin Fallavier, France) added in the medium. MICs of fosfomycin were also determined at pH 5 and 6. Each *in vitro* experiment was replicated at least in three independent experiments and the median values were reported for each strain.

### Mechanisms of fosfomycin resistance

Known mechanisms of fosfomycin resistance were determined for each strain exhibiting an MIC of fosfomycin ≥8 μg/ml as several of them have been found in strains with MICs from the susceptible range (9). Mutations in the genes involved in fosfomycin chromosomal resistance (i.e., *murA, glpT, uhpT, cyaA, ptsI, uhpA, uhpB, uhpC*) were determined by nucleotide sequencing after amplification by PCR, as previously described (9). The amino acid sequences were compared with those of *E. coli* CFT073 and K-12.

### *In vitro* bacterial growth rate

Growth rates at 37°C were measured in Luria-Bertani (LB) broth with various pH (5, 6, and 7) and in sterile-filtered pooled human male urine (pH=6), as previously described (9). For each strain and condition, maximal growth rate (MGR) and time to achieve MGR (Tmax) were measured in three independent experiments and the median values were reported for each strain.

### Murine pyelonephritis model

We used the ascending, unobstructed UTI mouse model previously developed by our group (26, 27). Eight-week-old immunocompetent CBA female mice (weight 20–23 g) were used. Bacterial inocula were obtained by overnight incubation in LB broth followed by centrifugation at 8000 × *g* for 15 min. Pellets were suspended in 1 mL of sterile saline solution to a final inoculum of 10^9^ CFU/ml. Pyelonephritis was induced after general anesthesia (with an intraperitoneal administration of 150 mg/kg of body weight of ketamine and 0.5 mg/kg xylazine) by injecting 50 μL (10^7^ CFU of *E. coli)* into the bladder through a urethral catheter. Urine was sampled for pH was determination in 5 mice just before bacterial inoculation and 48h after infection. For each strain, 18-24 mice were infected. Two days after inoculation, 8 to 14 mice were sacrificed before treatment (start-of-treatment controls), and 9-10 were treated over 24h by subcutaneous injections (fosfomycin 100 mg/kg q4h for 24h). In order to avoid unnecessary mice killing, we did not constitute a group of untreated mice (end-of-treatment controls) because previous study setting up the model showed that bacterial counts in kidney were stable for at least 5 to 10 days (26). Treated mice were sacrificed 24h after the last antibiotic injection to avoid the carry-over effect. Kidneys were aseptically removed and were homogenized in 1 mL of saline solution. Then, 100 μL of the solution or its dilution were spread onto MH agar plates and incubated at 37°C for 24 h. Selection of resistant mutants after *in vivo* exposure was sought by plating 100 μL of kidney homogenates onto MH agar containing fosfomycin at a concentration of 4 times the MIC for fosfomycin-susceptible strains and 2 times the MIC for fosfomycin-resistant strains. Kidneys were considered sterile if no colony grew on agar plate. In the absence of bacterial growth, the log_10_ CFU/g of kidney was calculated considering the growth of one colony and the weight of the kidney, as the method detection limit. This corresponded approximately to 1.8 log_10_ CFU/g of kidney. Assessment criteria for each strain were: i) bacterial load in kidneys expressed as log_10_ CFU/g; ii) the percentage of sterile kidneys.

### Fosfomycin dosing regimen

Single-dose plasma pharmacokinetic studies were performed on eight-week-old CBA female mice (weight 20–23 g) in order to determine the therapeutic regimen that best reproduced peak and trough plasma levels obtained in humans with a daily oral-dose of 3 grams of fosfomycin-tromethamine (20 and 5 μg/ml, respectively) (15, 18, 19). Dosing interval was chosen accordingly to reproduce area under the concentration-time curve (AUC) in the range of that obtained in humans with a single oral-dose of 3 grams of fosfomycin-tromethamine *(ie* up to 228 mg.h/liter, as compared with a range of 1400 to 1800 mg.h/L in human with the iv route) (13, 15, 18) since fosfomycin AUC/MIC ratio is the PK/PD index most closely linked to in vivo efficacy (24). This regimen was determined as 100 mg/kg every 4 h (15, 18, 19). Total drug concentrations were utilized in PK/PD analyses, as fosfomycin is not bound to plasma protein (15, 18).

### Fosfomycin sampling in plasma and kidney

Blood samples of at least 500 μL were obtained by intracardiac puncture from 4 anesthetized mice at 5 different intervals after a single subcutaneous injection of fosfomycin (100 mg/kg): 1h, 2h, 4h, 6h and 24h. After blood collection, plasma was separated by centrifugation. Kidneys were also removed at the same sampling times.

### Fosfomycin assays in plasma and kidneys

Concentrations of fosfomycin were determined using a triple quadrupole mass spectrometer, AquityÒ TQD (Waters, St Quentin en Yvelines), operated with negative electrospray ionization. Instrument parameters were optimized for fosfomycin (137 -> 79 m/z) and propylphosphonic acid (internal standard) (123 -> 79 m/z) transitions. Fosfomycin was extracted from samples (plasma and kidneys) via protein precipitation with acetonitrile. The chromatographic separation took place on an AcquityÒ UPLC BEH HILIC column with the dimensions 100 mm x 2.1 mm and 1.7 μm particle size (Waters, St Quentin en Yvelines). The lower limit of quantitation was 1 mg/L. Intraday and inter-day coefficients of variation in plasma were 8.5 and 11.1%, respectively, at concentrations ranging from 5 to 250 μg/ml. The possible influence of co-extracted matrix compound on detectability of target analyses was checked.

### Fosfomycin PK/PD analysis

For the PK analysis, concentration data of fosfomycin in plasma and kidney were separately fitted to 2-compartments models with extravascular administration (30), pooling data of all mice and using the ‘optim’ package of R statistical software (v3.4.0). For each fit, the pharmacokinetic model could thus be written as: 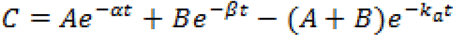. Parameters to be estimated were A and B, first and second macro-constants; α and β, first and second-rate constants, and k_a_, absorption rate constant. Data below the LLOQ were imputed to the LLOQ. We derived the AUC_0-24h_ in plasma (AUC_0-24h,plasma_) and kidney (AUC_0-24h,kidney_) and in in plasma to the time of sacrifice (AUC_0-44h_) from the model with a 100mg/kg administration of fosfomycin every 4 hours. Finally, AUC_0-24h_/MIC ratios were determined in plasma and kidneys for each strain used in the murine pyelonephritis model. We studied the relationship between the bacterial load in kidney and AUC_0-24h_/MIC ratios in plasma or kidney using linear regressions.

### Statistical analysis

MGRs of bacterial strains at pH 7, and studied the effect of pH on MGRs and time to achieve MGR using a Kruskall-Wallis non parametric test.

A 2-way analysis of variance was performed to study the effect of the infective strains and fosfomycin treatment on the bacterial loads in kidneys, and tested the interaction between the infective strain susceptibility and the treatment effect. We also compared the proportion of sterile kidney in fosfomycin-treated mice according to the infective strain using a Fisher exact test. Bacterial loads in kidneys from treated mice were compared between the 3 susceptible and the 3 resistant strains by the Mann and Whitney non parametric test. The relationship between the bacterial load in kidney and the AUC_0-24h_/MIC and AUC_0-44h_/MIC ratios in plasma and the AUC_0-24h_/MIC ratio in kidneys was studied using linear regression. Type-I error was set at 0.05, and two-tailed tests were used. All analyses were performed using R statistical software.

## Transparency declarations

Annabelle Pourbaix received Grants from the “Fondation pour la Recherche Médicale”, and from the “Société de Pathologie Infectieuse de Langue Française” for this work.

## Acknowledgements

We are indebted to Sara Dion and Louis Gary for technical assistance and Professor France Mentre for statistical assistance.

## Ethical approval

The pyelonephritis protocol (n° APAFIS#4950-2016021211417682 v4) was approved by the French Ministry of Research and by the ethical committee for animal experiment.

